# Applying Machine Learning to Investigate Long Term Insect-Plant Interactions Preserved on Digitized Herbarium Specimens

**DOI:** 10.1101/790899

**Authors:** E.K. Meineke, C. Tomasi, S. Yuan, K.M. Pryer

**Author notes:** Email addresses: EKM, CT, SY, KMP. Manuscript received October 1, 2019.

## Abstract

**Premise of the study:** Despite the economic importance of insect damage to plants, long-term data documenting changes in insect damage (‘herbivory’) and diversity are limited. Millions of pressed plant specimens are now available online for collecting big data on plant-insect interactions during the Anthropocene.

**Methods:** We initiated development of machine learning methods to automate extraction of herbivory data from herbarium specimens. We trained an insect damage detector and a damage type classifier on two distantly related plant species. We experimented with 1) classifying six types of herbivory and two control categories of undamaged leaf, and 2) detecting two of these damage categories for which several hundred annotations were available.

**Results:** Classification models identified the correct type of herbivory 81.5% of the time. The damage classifier was accurate for categories with at least one hundred test samples. We show anecdotally that the detector works well when asked to detect two types of damage.

**Discussion:** The classifier and detector together are a promising first step for the automation of herbivory data collection. We describe ongoing efforts to increase the accuracy of these models to allow other researchers to extract similar data and apply them to address a variety of biological hypotheses.

## Introduction

More than 350,000,000 pressed plant specimens are stored in the world’s 3,400 herbaria (Soltis, 2017). Collected over the past four centuries, they provide the most comprehensive view of Earth’s vegetation and how it has changed over time. In most cases, herbarium specimens are used to document plant biodiversity. However, they are now being applied to scientific efforts well beyond taxonomy and systematics, and in particular to global change biology. Millions of herbarium specimens were collected prior to the intensification of human influence on the planet, including the acceleration of climate change. The unique, long-term data preserved within herbarium collections can now help us understand the past and predict the future of global change (Heberling and Isaac, 2017; Lang et al., 2019; Meineke et al., 2018a; Meineke, et al., 2018b).

Until recently, the world’s herbarium specimens were under lock-and-key, accessible only to relatively few scientific specialists. Today, the digitization of herbaria is a global enterprise, and millions of high-resolution specimen images and their associated metadata are newly available in online public databases (Page et al., 2015; Soltis and Soltis, 2016). In their most widespread application outside of taxonomy and systematics, herbarium specimens have accelerated the study of plant phenological change. Reproductive structures such as flowers, buds, and fruits, along with the dates and locations associated with the specimens, can provide information on how phenological timing has shifted with climate (e.g., Davis et al., 2015). Notably, scientists have established that warmer temperatures are associated with widespread, earlier flowering times in temperate North America (Panchen et al., 2012; Primack et al., 2004). Though phenology has been a key focus of global change research using herbarium collections, it encompasses only a small amount of the long-term data that could be mined from digitized specimens.

In particular, herbaria are unmatched repositories for data documenting interactions between plants and associated species. Species interactions, such as those between plants and insects, are notoriously difficult to monitor over time, and data on species interactions spanning the Anthropocene are severely limited (Meineke and Davies, 2018). Plants have been engaged in a reciprocal war with the rasping, sucking, and chewing insects feeding on them for more than 400 million years. This long-term coevolutionary arms race between insects and plants is the basis for Ehrlich and Raven’s (1964) hypothesis that herbivorous insects have driven the evolution of plants, and in turn, that plant adaptations and defenses to insect attack have stimulated the diversification of insects. As such, relationships between plants and insect herbivores are central to ecology and evolutionary biology.

Herbarium specimens preserve signatures of insect damage (“herbivory”) through time on their leaves (Beaulieu et al., 2018; Meineke et al., 2018a). Herbivory encompasses diverse types of damage triggered by a wide range of insect taxa (for examples, see Fig. 1). Interestingly, similar signatures of insect damage have been documented on fossilized leaves. The paleobotanical community has already initiated critical studies that use and interpret these data preserved on fossilized leaves to understand changing species interactions over deep time between plants and insects (Wilf and Labandeira, 1999; Wilf et al., 2001). Herbarium specimens offer an opportunity to conduct parallel studies that analyze changing plant-insect associations over the period of anthropogenic change.

**Figure 1.**
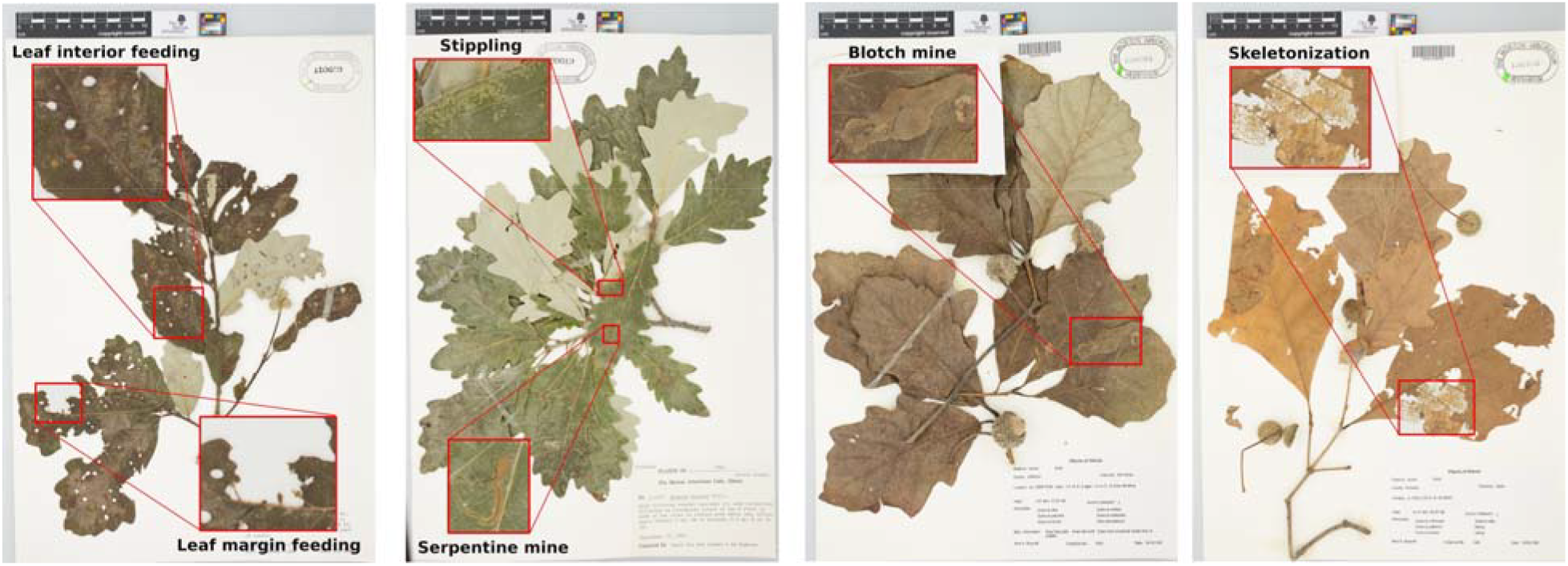
Herbarium specimens exhibiting a range of herbivory types made by different insect taxa, including examples of leaf interior feeding, leaf margin feeding, stippling, serpentine mines, blotch mines, and skeletonization. These are the types of herbivory for which recognition was automated in this study.

The only study, to date, to use herbarium specimens to quantify how herbivory has shifted over the past 100+ years was published by Meineke et al. (2018a), where over a period of two years, a single researcher painstakingly overlaid a physical grid of 5 ⨉ 5 cm cells on almost 600 herbarium specimens, developing novel methods for manually identifying and quantifying herbivory. They demonstrated that insect damage to four distantly related, woody angiosperm species in New England increased over the past 112 years by 23%, a pattern attributed to increased winter warming that promoted overwintering survival and/or range expansion of herbivorous insects. Meineke et al. (2018a) is one of the first studies to use data captured from herbarium specimens to investigate hypotheses about the ecological and evolutionary mechanisms driving herbivory. However, the amount of herbivory data that can become available for future study is constrained by how much data individual researchers can manually collect from physical herbarium specimens.

Here, we move beyond the manual procedures advanced by Meineke et al. (2018a) by developing novel automated methods to replace them, with the goal of allowing future studies on plant-insect species interactions to harness data from millions of herbarium specimen images available online. Similar to other studies that focus on plant reproductive parts and phenology (Lorieul et al., 2019), we develop machine learning algorithms that can recognize and quantify insect damage on digitized herbarium specimens. We then discuss ongoing improvements on these methods to facilitate the broadscale extraction of species interactions data.

## Methods

### Focal species

Meineke et al. (2018a) manually collected herbivory data from ca. 600 herbarium specimens of four woody angiosperm species distributed in eastern North America: *Quercus bicolor*, *Vaccinium angustifolium, Carya ovata*, and *Desmodium canadense.* We chose *Quercus bicolor* for this study because it exhibited the most diverse herbivory (i.e., the most types of herbivory per specimen). The other focal species we chose was *Onoclea sensibilis* from the fern lineage that is sister to seed plants because we wanted to test similar machine learning classifiers on plant species with diverse leaf morphologies. Both species are native to eastern and parts of central North America.

### A primer on machine learning

Machine learning involves the study and construction of computer algorithms that can learn and make predictions based on data. Predictors most relevant to our study are in the form of ***classifiers***, *i.e*., algorithms that predict the category in which some input data belong. For example, the input may be a rectangle within an image, and the output category may be one of several, pre-specified types of insect damage. To train a classifier, another algorithm called a ***trainer*** is given a ***training set*** consisting of many ***training samples*** (examples of inputs, together with the correctly annotated corresponding outputs). The trainer then adjusts the parameters of the classifier such that it will generate the correct output for as many of the training sample inputs as possible.

If the classifier has many parameters, the trainer can often fine-tune them so that the classifier’s answers are nearly perfect on the training set. However, the classifier will typically perform more poorly on previously unseen data. When this occurs, the classifier is said to have ***overfit*** the training set. In contrast, a classifier that does well even on new data is said to ***generalize*** well.

### Detection

Only the relatively small parts of a specimen image that contain insect damage are relevant to damage type classification, and a machine learning system must also learn how to detect these parts. Thus, an automatic analyzer has two stages, at least conceptually; the first stage is a ***detector***, and the second is a ***classifier***. The detector takes an entire specimen image as its input and outputs a collection of boxes that are likely to contain insect damage.

These boxes are similar to those that a human annotator would draw. The classifier then takes each of these boxes in turn and determines which type of damage it contains. The detector is a ***regressor***, rather than a classifier. This means that its output is not one of a small number of predefined categories, but rather an element out of a potentially infinite set (or at least an extremely large number) of options. Each output option for a detector is a list of image rectangles and each rectangle can be specified by the two coordinates of its upper-left corner and those of its lower-right corner. Thus, the detector outputs a list of quadruples of numbers, which is a much more complex object than a simple category out of a small set.

Detection is obviously an important task in machine learning, since it is rarely the case that an entire input image is relevant to the task at hand. Several detectors have been developed in the recent past (Erhan et al., 2014; Girshick 2015; Girshick et al., 2014; Ren et al., 2017). An interesting lesson learned from this research is that it is more effective and efficient to combine detection and classification into a single system that simultaneously finds boxes *and* determines the category for each of them (Hariharan et al., 2015; W. Liu et al., 2016; Redmon et al., 2016). Either way, detection solves a difficult problem because of the very large number of possible outputs and is, therefore, a data-hungry task. Table 1 shows that the experiments described in this paper are based on modest amounts of annotated data, and are therefore strongly colored by this complexity consideration, as will be discussed later on.

**Table 1.** Leaf damage categories annotated in our dataset. The last two columns denote “no damage” categories.

|  | Margin Feeding | Interior Feeding | Skeleton - ization | Stippling | Blotch Mines | Serpenti- ne Mines | Normal Margin | Normal Interior |
| --- | --- | --- | --- | --- | --- | --- | --- | --- |
| *Onoclea Sensibilis* | 39 | 28 | 8 | 0 | 11 | 11 | 46 | 25 |
| *Quercus Bicolor* | 456 | 616 | 215 | 184 | 28 | 18 | 206 | 223 |

### Herbivory annotation

We downloaded all of the high-resolution, digitized specimens available for *Quercus bicolor* and *Onoclea sensibilis* from the SouthEast Regional Network of Expertise and Collections National Science Foundation Thematic Collections Network (SERNEC: http://sernecportal.org/portal/; Appendix 1). We then manually annotated 109 images of *Q. bicolor* and 15 images of *O. sensibilis* specimens for all instances of clear insect damage. We used the VGG Image Annotator v. 2.0.4 (Dutta and Zissermann 2016; Dutta 2019) to draw bounding boxes and assigned a damage category to each box. Table 1 shows the number of instances annotated for each of the six damage types we investigated in this study (leaf margin feeding, leaf interior feeding, skeletonization, stippling, blotch mines, and serpentine mines). The categories “normal margin” and “normal interior” represent boxes drawn on undamaged parts of the specimens and constitute “negative examples” for the training set. For an example of each annotation category, see Fig. 1.

### Detection and Classification Experiments

High-quality automatic detection and classification of leaf-damage boxes require more annotated data than we currently have. Therefore, we split the detection and classification experiments into two groups and made simplifications in each of them. Specifically, in the first group of experiments we detect and simultaneously classify only two types of damage, namely, leaf margin feeding and leaf interior feeding, for each of which the number of available annotations is in the several hundred (Table 1). In the second group of experiments, we classify each of a set of manually annotated boxes into one out of all eight categories (six damage categories and two “no damage” categories, see Table 1). For both experiments, we extracted a subset of the data to be used for training, and we use the rest for performance evaluation. This data split is described in more detail below for each experiment.

#### i. Data Split for Detection Experiments

Of the 120 images that were manually annotated, we retained the 105 images that contained some margin feeding and interior feeding annotations. Of these images, we selected 83 at random for training and kept the remaining 22 for testing. The detection methods we used in our experiments work best with non-overlapping annotation rectangles. Because of this, we discarded a small fraction of annotations when overlaps occurred. In the end, training images contained 348 instances of margin feeding and 444 of interior feeding. Test images contained 106 instances of margin feeding and 124 of interior feeding.

To sharpen the detector’s ability to distinguish between these types of damage and normal parts of a leaf, or from other types of damage, we also provided what are called *hard negative examples* to the detector. Specifically, we added a third category, which can be interpreted as a “null” category (that is, neither interior feeding nor margin feeding) and placed in it all the boxes in the training images that were annotated as different types of damage by the human annotators. The detector was then trained to detect boxes of the three types, but all detections from the null category were eventually ignored during testing. This mechanism allows the detector to learn a more detailed understanding of the boundary between, *e.g.*, an instance of margin feeding and an instance of a normal margin. Intuitively, we are showing the detector not only what margin feeding looks like, but also what it does <u>not</u> look like. This strategy has been found useful in a variety of computer vision systems (Dong et al., 2017).

#### ii. Data Split for Classification Experiments

Annotation boxes are rectangles of arbitrary sizes that are drawn to enclose the leaf damage as tightly as possible. We used deep neural networks, described in more detail later, as classifiers. Because a typical deep neural network expects an input box of a fixed size, we wrote software that samples squares of 224 × 224 pixels from each of the annotation boxes. This is the smallest image size expected by a wide range of current deep-learning architectures. For annotation boxes smaller than this size, a 224 × 224 square was extracted from the original image, with the annotation box at its center. For larger annotation boxes, several 224 × 224 squares were sampled, providing a crude form of data augmentation. For the margin feeding, interior feeding, and normal margin categories, it was important that each box contains both leaf and background. To this end, we wrote image segmentation software to identify the boundaries between leaf and background, and only sampled boxes for which the leaf covered at least 40 percent of the area. Our sampling procedure resulted in a dataset with 6616 samples that were split uniformly at random into 4157 samples for training, 1350 for validation, and 1109 for testing. This split was executed at the annotation-box level so that all boxes from a given annotation fall into exactly one of the three sets. A classifier was trained on training samples and evaluated on test samples. Validation samples are used during training to estimate when the classification algorithm has reached its best generalization performance.

### Experimental Machine Learning Architectures and Pre-Training

#### Detection Network Architecture

We used the Single Shot Detector (SSD) with a VGG16 base classification network (Liu et al., 2016) to simultaneously detect and classify interior feeding and margin feeding. This detector makes a very large number of box hypotheses in the input image. Box hypotheses differ by both their location in the image and their shape and size. The base classification network then classifies each box hypothesis and computes a score for each of them, which measures the network’s confidence in the classification result. Each high-score box is output as a detection, together with the category that yielded the maximum score for that box. A box is not output if it has a score that is high, but lower than that of another box with which it overlaps. This criterion prevents overlapping detections to be output.

#### Classification Network Architecture

The damage type classifier was adapted from a Residual Net architecture (He et al., 2016) with 18 layers. This is a standard, small deep network that has shown good performance in a variety of image recognition tasks. Originally designed to distinguish among 1000 object categories, we adapted it by changing its last layer to work with our eight damage type categories instead. Details of the classifier’s architecture are given in Appendix 2.

#### Pre-Training

The networks we use in our experiments have large numbers of parameters to be estimated during training. This necessitates large amounts of annotated data for training. In order to address the disparity between the size of our training set and the number of parameters to train, we used what is called “pre-training” in the literature. Specifically, we started our training with neural networks (both for the base classifier in the SSD detector and for the damage type classifier) that had already been trained on a classification task that is entirely different from the target task and for which ample data are available, namely, the ImageNet database (Russakovsky et al., 2015). This dataset contains millions of labeled images of ordinary scenes and objects in thousands of categories. This initial phase is called the pre-training phase, and its purpose is to replace random values for the network parameters with values that at least relate to plausible images. We then further trained the networks on our annotated images. This technique, which is commonly used in computer vision, has also proven useful in the domain of plant species identification (Carranza-Rojas et al., 2017).

## Results

### Damage Detection

Simultaneous detection and classification, as performed by the SSD (Liu and Stiling, 2006) or similar detectors (Redmon et al., 2016; Sermanet et al., 2013), requires large amounts of annotated images per class. In our data, only interior feeding and margin feeding damage have several hundred manually annotated boxes, and after the data split these two categories have 392 and 321 training examples, respectively. These numbers are barely enough for detection classification and, even so, we had to use both pre-training and hard negative examples, as described earlier, in order to achieve acceptable performance. Because of this, we limit our detection experiments to these two categories, as explained above.

#### Detailed Results

Figures 2 through 4 show *all* the experimental results for the 10 images we ask the annotator to exhaustively annotate all the margin feeding and interior feeding boxes. Specifically, Fig. 2 shows all true-positive detections of interior damage. These are instances where human annotator and detection algorithm agree, in the sense that their two boxes overlap at least 50 percent of the size of the smaller of them. The left collage (Fig. 2) shows results for interior damage (green boxes; dashed line for annotations and solid line for predictions) and the right for margin damage (red boxes; dashed line for annotations and solid line for predictions). Looking at the example in the top left box in the collage on the left of Fig. 2, we see that a human annotator may annotate an entire area of sparse damage with a single, large box, while the algorithm may detect small separate sub-areas as distinct instances or, in this case particularly, miss several of the sub-areas altogether. This is also the case for margin damage (collage on the right of Fig. 2).

**Figure 2.**
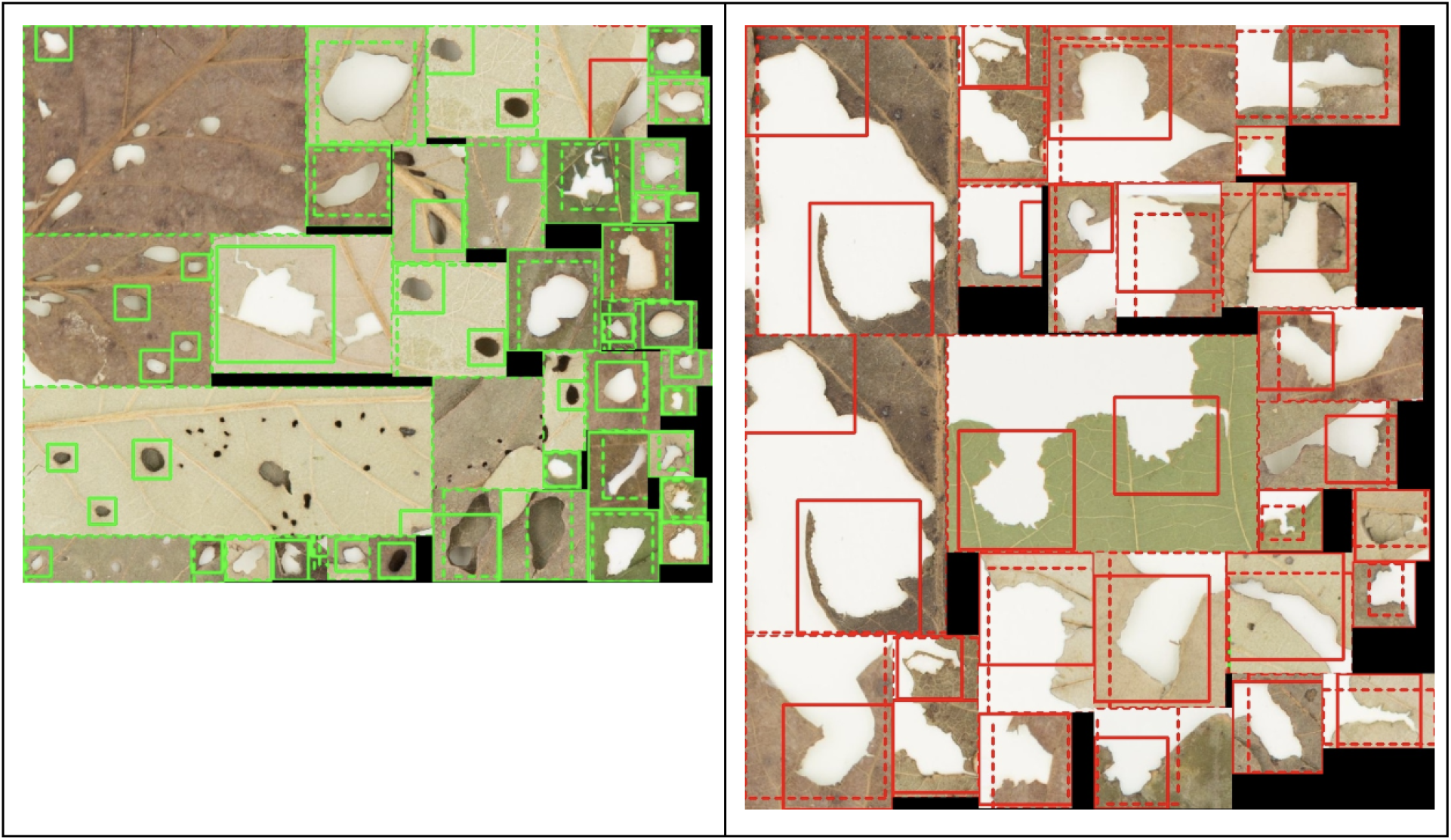
A collage of all true-positive detections for interior leaf damage (left) and leaf margin damage (right) in 22 test images. A true positive is an instance of damage that was annotated by a human ***and*** detected by the detector. In these images, dashed boxes are human annotations, and solid boxes are detector results. Green boxes are for interior feeding, red ones for margin feeding.

Figure 3 shows all false negatives, that is, all the human annotation boxes that the detector missed. Comparison of the two left collages in Figs. 2 and 3 suggests that the detector “got used” to identifying interior damage as being more or less oval holes and did not have enough examples of more complex shapes to adapt to these during training. Perusal of both right collages in Figs. 2 and 3 suggests that the shapes of margin feeding instances are very varied, and many more examples would be necessary for the network to learn an appropriate model of them.

**Figure 3.**
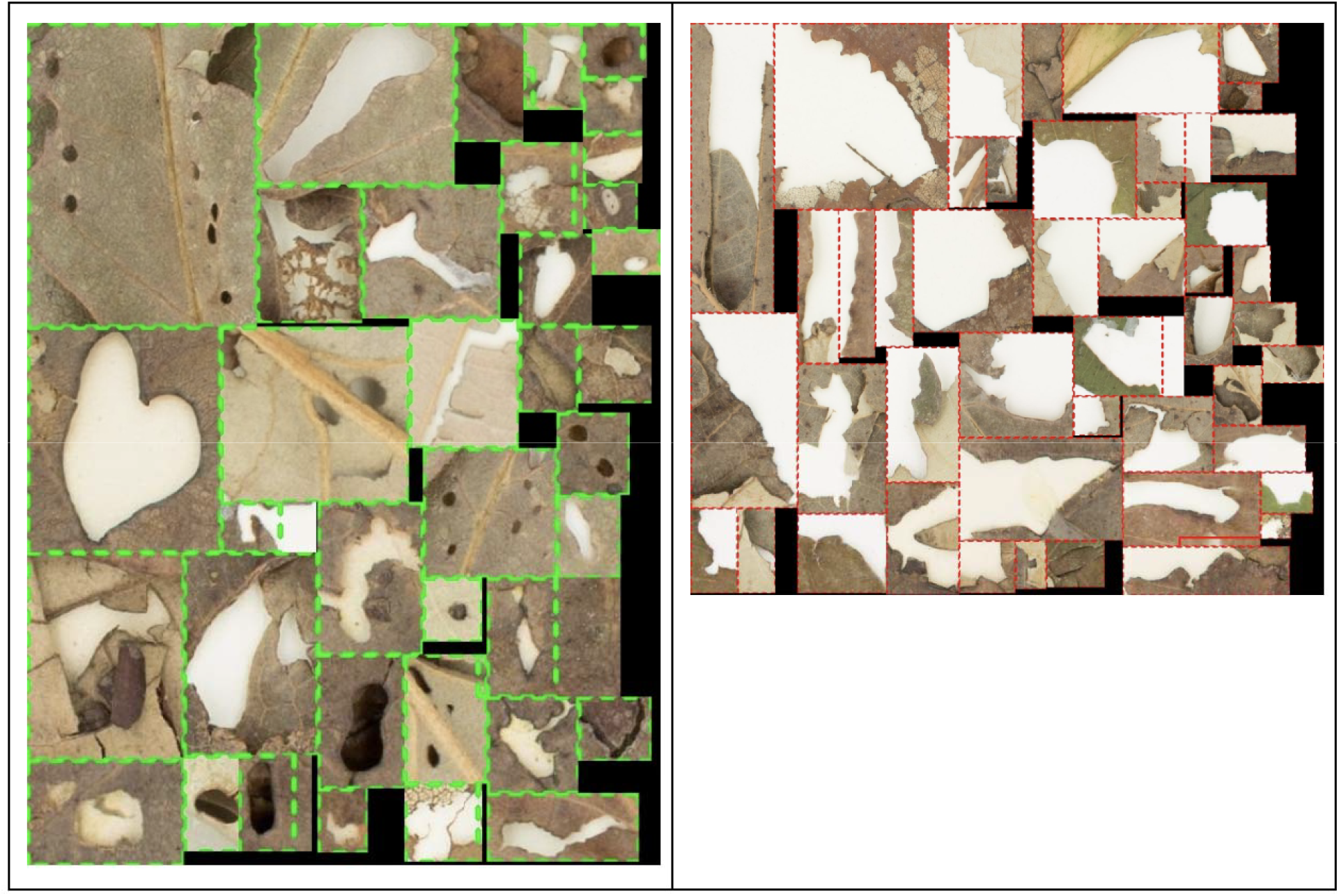
A collage of all false-negative detections for interior leaf damage (left) and leaf margin damage (right) in 22 test images. A false negative is an instance of damage that was annotated by a human but was missed by the detector. See caption for Fig. 2 for the meaning of line styles and colors.

Figure 4 shows all false positives, that is, image regions that the detector thought to be examples of interior (left) or margin (right) feeding, but that the human annotator did not mark. First, there are rather few such boxes in these two collages, that is, the false-positive rate is low. Second, some of the detections seem genuine damage, although not necessarily of the declared type. For example, many of the oval lesions seen in this figure could be from disease (perhaps fungal) rather than insects. An additional consideration about Fig. 4 is the misclassification of digits from the printed text in the specimen images as either interior (left) or margin (right) feeding. It would be relatively straightforward to develop image pre-processing routines that exclude these areas from consideration. We left these detections in the figure because they clearly point to over-fitting; the digits on the left all contain closed ovals, and this reinforces the observation we made for Fig. 3 (left), that the detector takes anything that looks like an oval and classifies it as interior damage. Similarly, all digits on the right contain at least some open digits, whose shape could be interpreted as the profile of part of a leaf’s boundary. In this context, over-fitting means that the detector formed a naive model of what these two types of damage look like, based on sparse data. More data would be needed to develop a more apt model.

**Figure 4.**
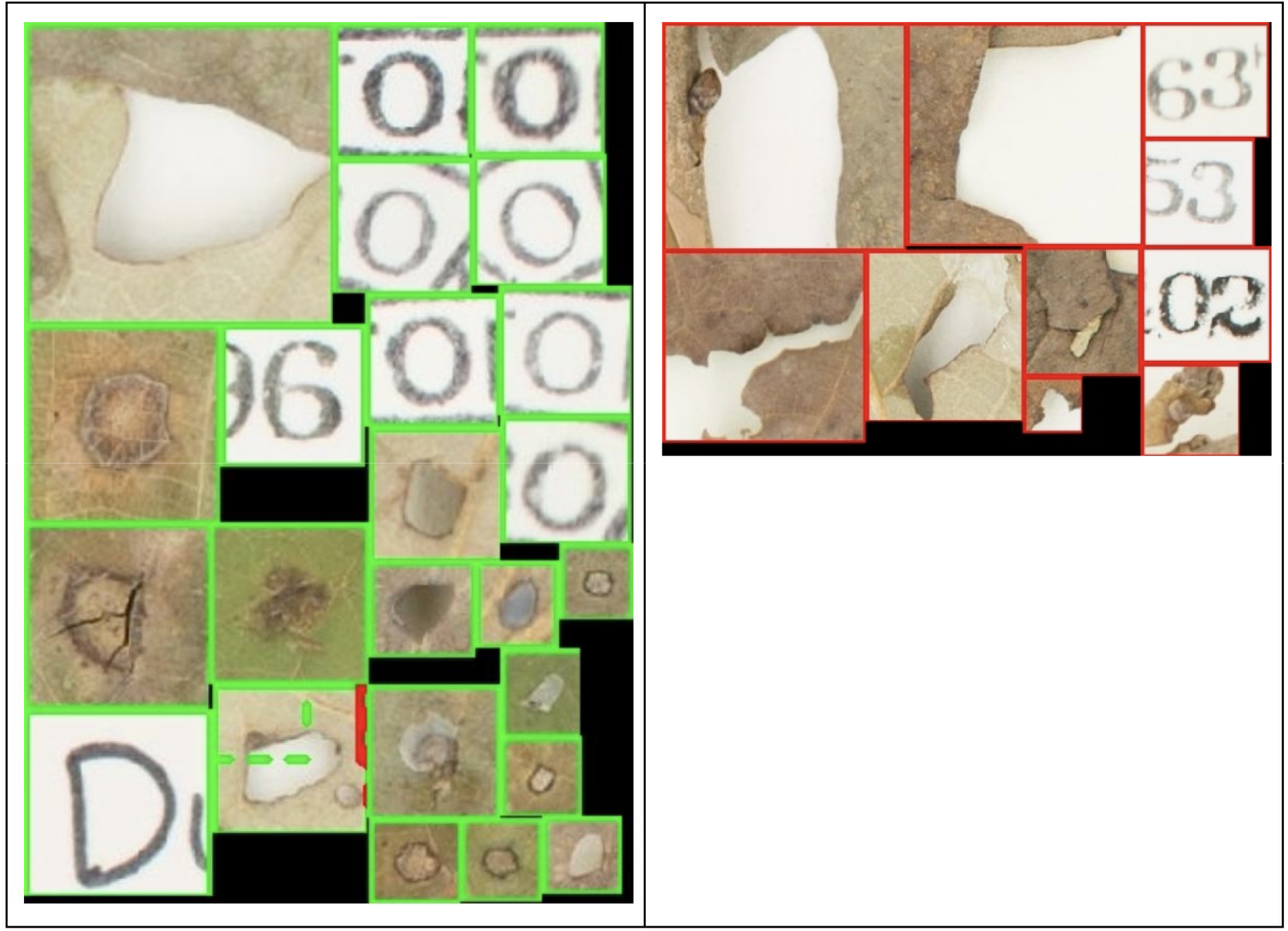
A collage of all false-positive detections for interior leaf damage (left) and leaf margin damage (right) in 22 test images. A false positive is an image region that was detected by the algorithm but not marked as either interior feeding or margin feeding by the human annotator. See the caption of Fig. 2 for the meaning of line styles and colors.

#### Issues with Aggregate Measures of Performance

It is standard practice in machine learning papers to give aggregate measures of performance such as overall error rates or accuracy. For a detector, one very popular measure is the average *Intersection over Union (IoU)*: A human annotation box H and a detected box D are said to match if they overlap, and the extent of overlap is measured by the area of the overlap region (the intersection) divided by the area of the union of the two regions. If H and D are identical, this measure is equal to 1, and if H and D do not overlap the measure is 0. The average IoU over all boxes in the test set then gives an overall measure of performance.

For damage detection, this measure would be very misleading, and the examples discussed above show why. What constitutes “a single instance” of damage is a poorly defined concept, and even when the human and the detector happen to disagree, both results are sometimes plausible. For instance, even if the detector had found all the small holes in the top-left box of Fig. 2 (left), the intersection would be the aggregate area of the small holes, and the union would be the area of the human annotation. The ratio of these two quantities is small, suggesting poor performance. However, most people would agree that either interpretation (one large box or several small ones) effectively identifies the same damage. Similar considerations hold for other aggregate measures of performance, which we therefore do not compute.

### Damage Classification

Our damage classifier network returned the correct answer in 81.5% of the 1109 testing samples (those that were <u>not</u> used for training or validation). The confusion matrix for our results is shown in Table 2. Each row in the table corresponds to a true answer, and each column corresponds to the answer given by the classifier. Each entry in the table is the number of times an instance from a row category was classified into a column category. For instance, the entry 39 in the first column means that a normal margin was mistaken for an instance of margin feeding in 39 cases. An ideal confusion matrix would have non-zero entries only on the diagonal, which contains the number of correct classifications (bold and highlighted in Table 2).

**Table 2.** Confusion matrix with correct / incorrect predictions made by our classifier on our dataset.

|  | Margin Feeding | Interior Feeding | Skeleton - ization | Stippling | Blotch Mines | Serpentine Mines | Normal Margin | Normal Interior |
| --- | --- | --- | --- | --- | --- | --- | --- | --- |
| Margin Feeding | **114** | 13 | 0 | 1 | 0 | 0 | 14 | 0 |
| Interior Feeding | 4 | **128** | 2 | 0 | 0 | 0 | 0 | 0 |
| Skeletonization | 1 | 1 | **31** | 2 | 2 | 0 | 0 | 0 |
| Stippling | 1 | 0 | 3 | **30** | 2 | 0 | 0 | 1 |
| Blotch Mines | 5 | 3 | 81 | 13 | **40** | 0 | 0 | 1 |
| Serpentine Mines | 0 | 0 | 1 | 0 | 1 | **3** | 0 | 1 |
| Normal Margin | 39 | 0 | 0 | 0 | 0 | 0 | **334** | 1 |
| Normal Interior | 1 | 0 | 0 | 1 | 0 | 0 | 6 | **224** |

The damage classifier was relatively accurate for those categories with at least one hundred test samples (margin feeding, interior feeding, normal margin, normal interior, see Table 2; and therefore at least about 400 training samples since the training set is about four times as large as the test set). An exception was some confusion between normal margin and margin feeding (39 in the first column and 14 in the next-to-last column). The results for more sparsely represented categories are not very meaningful statistically except for blotch mines that are very often confused with skeletonization damage (81 cases). Interestingly, the damage that insects make within blotch mines is skeletonization, so the error is not overly surprising and indicates to us that critical refinements will be needed in specifying our damage types moving forward.

## Discussion

We developed automated techniques that detect insect damage for two categories (leaf interior feeding and leaf margin feeding) and categorize a diverse array of insect herbivory by damage morphology from pre-segmented damage boxes. To the best of our knowledge, this is the first attempt to develop automated techniques that will scan pressed plant specimens and identify clear instances of insect damage. The overall performance observed in this study including only a modest dataset is promising. Specifically, results of detection are mixed (Figs. 2-4), while herbivory classification results are objectively promising, with an accuracy of 81.5%. Future studies can improve the accuracy of the tools developed here.

The detection algorithm performed qualitatively well at detecting damage where a human annotator detected damage. This is a promising initial step in the development of a model that can locate specific types of damage. However, for the detector, it is clear that much more data will be needed for performance that is accurate enough to be used to extract data to test ecological and evolutionary hypotheses. For both detector and classifier, the confusion between damaged and normal margins speaks to the need for a damage mask to focus the damage classifier’s attention on the margin itself. The confusion between blotch mines and skeletonization indicates that texture and color descriptors, perhaps in the form of correlograms, will be needed to help make more nuanced distinctions. We did not use either damage masks or correlograms in our preliminary experiments.

We knew that one of the key challenges of this project would be to annotate enough instances of different types of leaf damage to accurately train the damage detector and the damage classifier. The size of the training set is critical. As other researchers have discovered—even in the domain of leaf classification—thousands of images are needed for the simpler task of binary classification of an entire specimen image into mercury-stained or unstained (Schuettpelz et al., 2017), and hundreds of thousands are needed for more complex tasks, such as species classification (Carranza-Rojas et al., 2017). As discussed above, box detection is even harder. We expect that tens of thousands of annotations per damage type will eventually be necessary for high quality results, and it will take time and effort to accomplish this in follow-up studies.

Anecdotally, it can be assumed that botanists may prefer to collect specimens with little or no insect damage. For this reason, any herbivory quantified on specimens is expected to be a conservative estimate of total herbivory experienced by plants. Thus, it is important to acknowledge and/or account for these biases as machine learning methods continue to be developed. Importantly, the tool we present could be used on any pressed leaves, opening the possibility for automated scoring of percent leaf herbivory in situations where this down-bias is not an issue. For example, our techniques could be applied to leaves from field or greenhouse studies, which is perhaps a more common application in ecology than measuring herbivory on herbarium specimens (e.g., Johnson et al., 2016; Turcotte et al., 2014).

Paleobotanists have used insect damage data from fossils to quantify changing patterns of insect diversity during periods of warming in the fossil record (Labandeira et al., 2002; Wilf and Labandeira, 1999; Wilf et al., 2006). We show here that not only could herbaria offer a parallel resource for examining modern changes in herbivory, but also that we may be able to use automated techniques to extract damage types from specimens, allowing for the possibility of “big data” extraction. This is important because some limited data that are available on insect abundances (Boyle et al., 2019; Wepprich et al., 2019) and biomass (Hallmann et al., 2017; Lister and Garcia, 2018) over time suggest that insects are in decline in the Anthropocene. However, these studies are highly debated today (Ries et al., 2019; Wepprich, 2019; Willig et al., 2019) and biomass studies often represent data collected from only two timepoints, one before and one after the acceleration of climate change. If the automated techniques described here are developed further and harnessed to their full potential, herbaria will offer an unprecedented opportunity to assess changing insect damage and diversity across broad scales of space, time, and plant phylogeny.

## Appendices

**Appendix 1** is provided as a. xlsx file.

### Appendix 2 Classifier and Training Details

Our neural net had an input convolutional layer with a 7 by 7 kernel, stride 2, batch normalization, a Rectified Linear Unit (ReLU), and a max-pooling layer with a 3 by 3 kernel and stride 2. Four standard residual blocks followed, each with four convolutional layers with 3 by 3 kernels, batch normalization, and ReLU. After a 1 by 1 convolution to adjust the output size to a vector of length 512, a final, fully connected layer changed the number of outputs to 8, the number of categories in our experiments. Classification was achieved by identifying the largest entry in a soft-max transformation of the output. The exact architecture can be downloaded and modified with the following PyTorch commands:

~~~
model = torchvision.models.resnet18(pretrained=True)
model.fc = torch.nn.Linear(model.fc.in_features, 8)
~~~

The model was trained with stochastic gradient descent with a constant learning rate of 0.0002 and a momentum of 0.9, and the cross entropy loss was used as the risk function to minimize.

## Acknowledgments

This work was supported by a Duke University Arts and Sciences Council Faculty Research Grant to KMP. We would like to thank Conrad Labandeira for guidance on scoring herbivory categories, and iDigBio along with its various Thematic Collections Networks for making herbarium specimens digitally available for study.

## Author Contributions

EKM, CT, and KMP designed the study, performed analyses, and wrote the paper. CT and SY designed and implemented machine learning approaches.

## Data Accessibility Statement

All data are published in the Appendix.

